# Characterization of BSN175: A Drug to Treat Prader-Willi Syndrome

**DOI:** 10.1101/448555

**Authors:** Alyson Ackerman, Robert A. Lodder

## Abstract

The purpose of this research is to choose the best analytical method for determining stability of BSN175. This research uses an accelerated stability study to compare the decomposed and stable drug using IR and ^1^H NMR spectroscopy. The Bootstrap Error-adjusted Single-sample Technique (BEST) was used to compare the effectiveness of these analytical methods and choose the best option for determining stability.

## INTRODUCTION

Prader-Willi syndrome (PWS) is a condition that affects thousands of children worldwide. Characterized by weakness in infancy and obesity, behavioral issues, and an insatiable appetite from around the age of two onwards^1^, this condition is all-consuming and can be devastating to those experiencing it and their families. One of the most recognizable symptoms is extreme obesity, with 50% of patients exceeding normative percentile ranges (defined as 3-97^th^ percentiles). For males aged 18, 25% exceeded 118 kg (260 lbs) and 25% of females aged 18 exceeded 102 kg (225 lbs).^2^ Type II diabetes frequently accompanies this obesity. While there is no cure for Prader-Willi, there are ways to manage the symptoms. Sugar substitutes are used to reduce caloric intake and sate those who cannot control their appetites. BSN175 is one such that is being investigated for use in treatment of PWS.

BSN175 is a carbohydrate drug. A major benefit of BSN175 is that while it is sweet like table sugar, it is not metabolized in the same way, and has a much lower calorie content along with low glycemic index. (The glycemic index is a measure of how much a sugar raises a person’s blood glucose level following consumption.) While glucose, sucrose, and fructose have glycemic indices of 100, 68, and 24, respectively; BSN175 has a glycemic index of 3. Additionally, BSN175 promotes glucokinase activity, which increases the rate of glucose to glycogen transfer. This reduces blood glucose levels.^3^ Therefore, this is an ideal form of treatment for PraderWilli as it satisfies patients’ appetites as well as reduces their serum glucose levels. It is especially desirable as a form of treatment as it can be incorporated into dosage forms resembling food products. This will aid in patient compliance with taking the medicine, especially as many patients are children.

To provide quantifiable, reproducible analysis of BSN175, the active pharmaceutical ingredient needs to be easily identified and differentiated from other possible impurities. Several spectroscopic and other analytical methods will need to be applied to the drug to build a package that can be used to determine the identity and purity of D-tagatose to meet FDA regulations for quality and safety.

This study focuses specifically on stability of the drug. An accelerated stability study was performed on Dtagatose and the stable and decomposed BSN175 were ^1^H both analyzed with NMR and infrared (IR) spectroscopy.

Following the collection of data, the Bootstrap Error-adjusted Single-sample Technique (BEST) was implemented to determine which method was preferred for determining stability. The BEST method calculates the distance between data clusters in multidimensional standard deviations (MSD).^4^ It is similar in nature to the Mahalanobis distance, but does not require that the number of rows exceed the number of columns. Because the BEST algorithm is not dependent on this matrix shape requirement for matrix factorization, it is suited to many more applications. The results of this statistical analysis will reveal which analytical test is the best choice for determining stability.

## METHODS

### Decomposition of BSN175

Approximately 5 grams of the BSN175 (Biospherics.net, Betrhesda, MD) was placed in a clean, glass dish. This dish was placed in an oven. The initial temperature was 52 degrees C. The temperature was increased 11 degrees C every ten minutes until the white BSN175 powder turned yellow.

The oven was 219 degrees C when the dish was removed. The BSN175 was cooled to room temperature, then it was scraped off of the glass dish and reground into a powder with particles that matched the size of the stable BSN175.

### IR Analysis

Both the stable and decomposed BSN175 were ground using mortar and pestle into a fine powder. Six samples of each were analyzed using a Thermo-Scientific Nicolet iS50 FT-IR.

### NMR Analysis

Approximately 5 mg of the BSN175 was added to ∼2 ml of D2O in a 400 mHz NMR tube. The tube was placed into a Jeol NMR and the sample was processed with 16 scans from −2 to 16 ppm. Six samples each of the decomposed and stable drug were analyzed in this way.

### Statistical Analysis

The NMR and IR data collected were arranged into matrices using MATLAB software. The BEST programs were used to determine the deviations between the center of the stable data and each set of decomposed data sets for NMR and IR results. A control was determined by using the same deviation command on only the stable set of data, measuring each individual data set’s deviation from the center of all stable data for both NMR and IR. The entire code used for this analysis can be found in Appendix I. This includes the two programs and commands implemented on the four types of data sets (each containing six trials): stable IR, decomposed IR, stable NMR, and decomposed NMR. The numerical results that can be found in Table 3 can also be found in this command section in bold font.

## RESULTS/DISCUSSION

Following the collection of data, each set of analytical spectra were combined onto the same plot for comparison. The infrared spectrum (Figure 1) contains 12 total sets of data: 6 of the stable BSN175 and 6 of the decomposed BSN175. Visually, these combined spectra show how precise the data for both the decomposed and stable BSN175 are, with little deviation between the sets of data for each form of the BSN175. The primary stretch of note is around 3400 cm^−1^, indicating the presence of an – OH group, as expected. This stretch is much sharper for the stable than decomposed BSN175, indicating some non-hydrogen bonding −OH groups. In general, the decomposed spectra are smoother, with broader stretches and less defined peaks. Of note, there is a small additional peak present in the decomposed spectra at 1645 cm^−1^, which is characteristic of an alkene stretch, though usually much larger and sharper. This information could aid in determining how this BSN175 decomposes and what compounds are produced. Currently, this information is largely unknown about caramels in general.^5^

**Figure 1.**
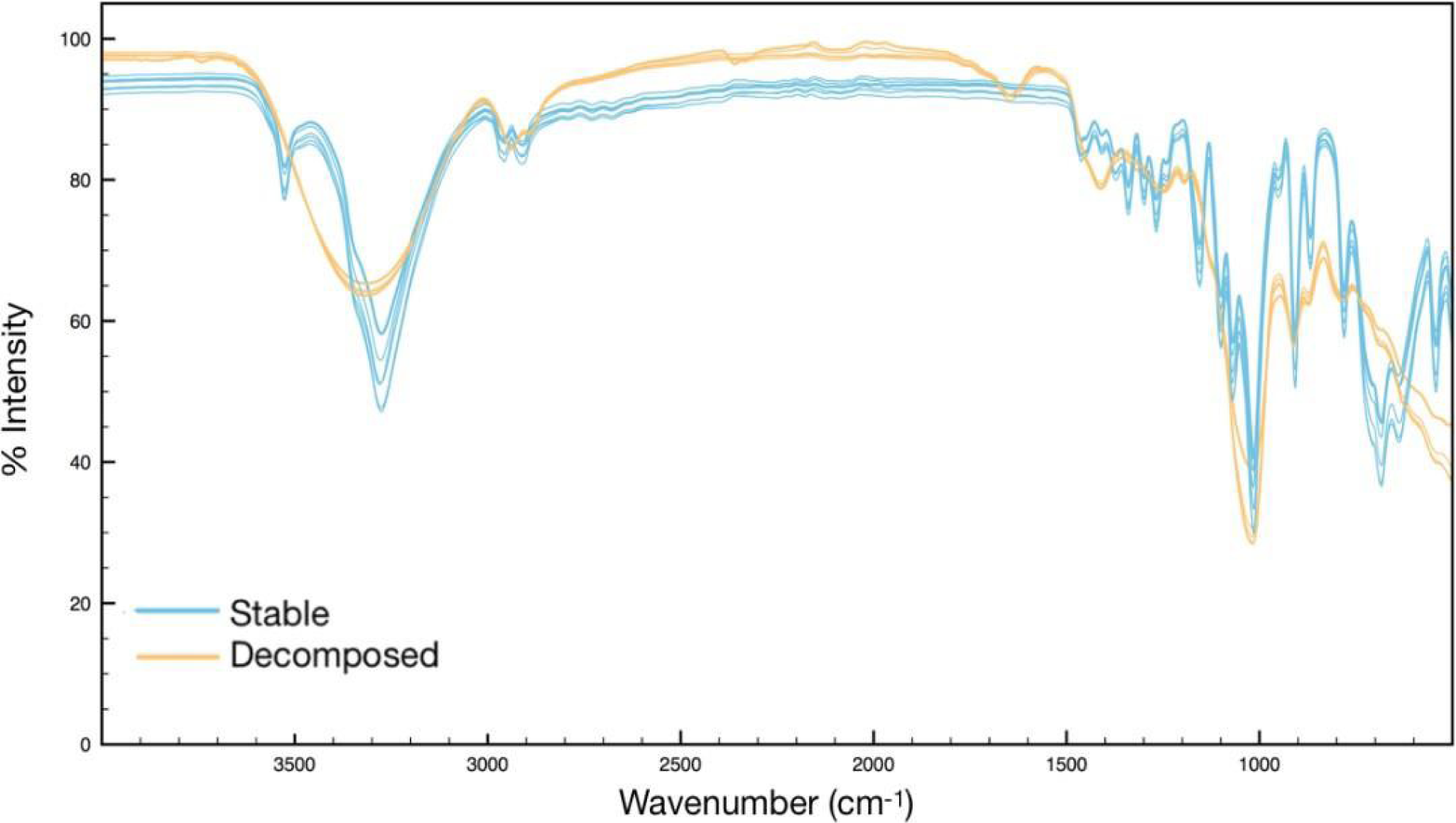
Infrared spectra of stable and decomposed BSN175

Stable and decomposed forms of the drug were then analyzed using ^1^H NMR (Figure 2). The results of the ^1^H NMR spectrum are analyzed below in Table 1 and the hydrogens assigned to the various peaks from the spectrum.

**Figure 2.**
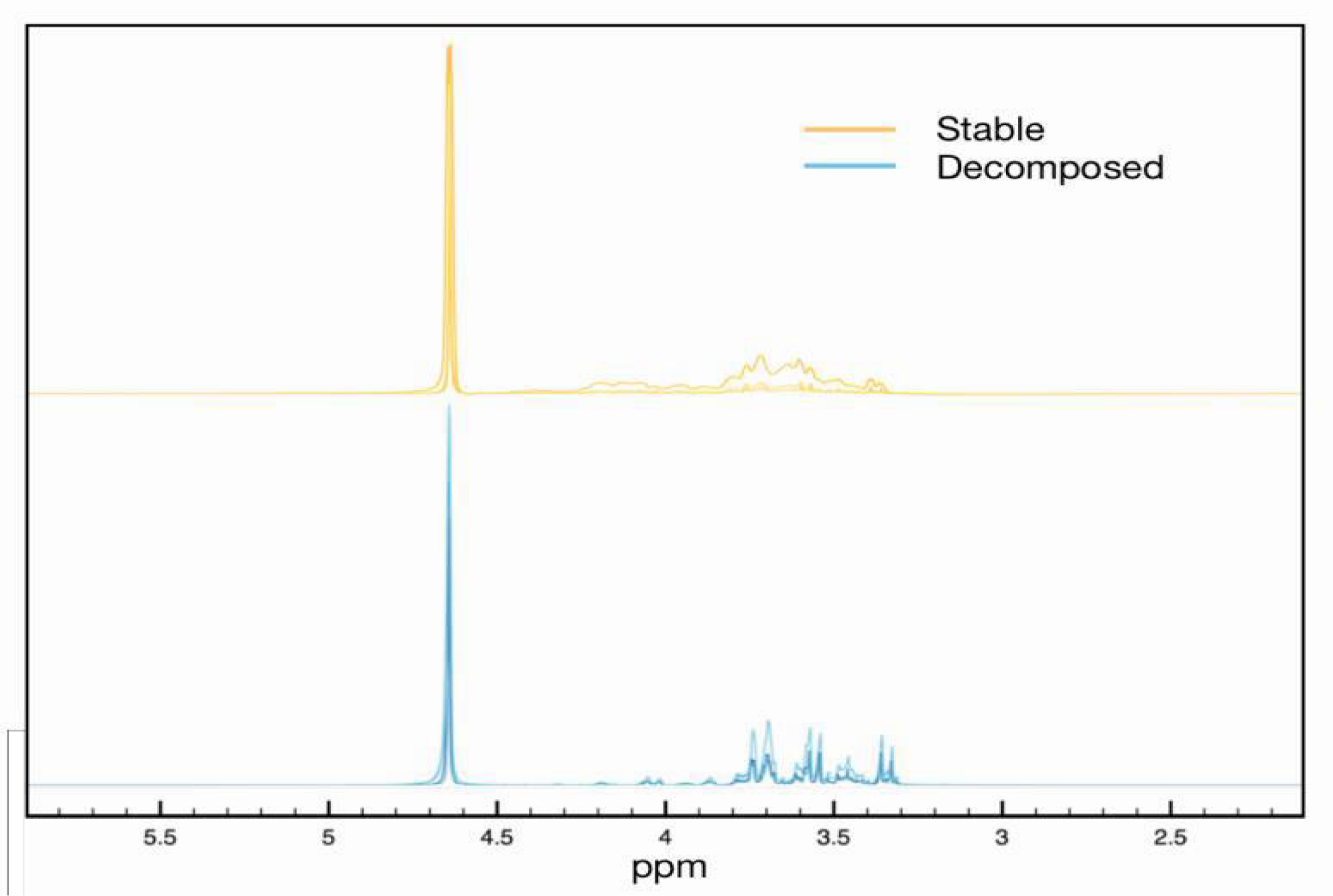
^1^H spectra of stable and decomposed BSN175

**Table 1.** ^1^H NMR analysis of shift, multiplicity, and integration for BSN175

| Proton | Chemical Shift (ppm) | Multiplicity | Integration |
| --- | --- | --- | --- |
| a | 4.653 | s | 4 |
| b | 3.741 | d | 1 |
| c | 3.584 | dd | 1 |
| d | 3.374 | dt | 1 |

There are four clear clusters on the spectra, which align to the four types of hydrogens present in the molecule. However, there are also some smaller clusters present, the largest of those presenting at 3.481 ppm. Further characterization is desirable to determine the source of this. The data for all six samples of stable and decomposed BSN175 (a total of 24 sets) was then submitted to MATLAB and the BEST statistical analysis was performed (see Appendix I for code). The multidimensional distances in standard deviations produced by the program can be found in Table 2 below. Note that the stable/stable deviations are a measure of one stable sample’s data from the center of all stable samples’s data combined and act as a control. The stable/decomposed deviations are a measure of one decomposed sample’s data from the center of all stable samples’s data combined.

**Table 2.** Summary of BEST statistical analysis results

|  | NMR |  | IR |  |
| --- | --- | --- | --- | --- |
|  | Stable/Stable | Stable/Decomp. | Stable/Stable | Stable/Decomp. |
| **Trial 1** | 1.25 | 3.50 | 1.41 | 35.05 |
| **Trial 2** | 1.50 | 3.63 | 0.78 | 24.74 |
| **Trial 3** | 0.80 | 1.24 | 1.34 | 23.46 |
| **Trial 4** | 0.74 | 1.19 | 1.44 | 22.11 |
| **Trial 5** | 1.07 | 4.28 | 1.42 | 21.72 |
| **Trial 6** | 1.03 | 4.31 | 1.73 | 25.71 |
| **AVERAGE** | 1.06 | 3.02 | 1.35 | 25.46 |

The stable/decomposed deviation are larger than the stable/stable (or control) deviations for both analytical methods, indicating that there is a quantifiable difference between the stable and decomposed drug.

The greater the distance in multidimensional SDs between the stable and decomposed drug, the easier it is to determine if the BSN175 is decomposing. Therefore, IR spectroscopy (25.46) is superior to NMR (3.02) for determining the thermal stability of BSN175. This can also be determined qualitatively by observing the IR and NMR spectra above in Figures 2 and 3. The visual difference between stable and decomposed BSN175 is much easier to see in the IR spectrum than the NMR spectrum, confirming the numerical data. However, in practice, using the BEST analysis with IR would provide a certain answer for whether the BSN175 is decomposed without the estimation required for visual determination.

## CONCLUSION

The results of this experiment indicate that IR spectroscopy is the best choice for routine monitoring of the stability of BSN175. The spectra produced for the stable and decomposed drug show a visually clear difference between the two. Additionally, the BEST standard deviation average of 25.46 is sufficiently large that determining the condition of the drug is not left up to interpretation, especially in comparison to the average stable-stable deviation of 1.35. The results from the ^1^H NMR are not as definite in determining stability. With a stable-decomposed deviation of 3.02 and stable-stable deviation of 1.06, there is quantifiable difference between the stable and decomposed BSN175. However, two of the decomposed samples tested provided deviations of only 1.240 and 1.187, which are both lower than two of the stable-stable deviations. This suggests that the ^1^H NMR spectrometry would not be as reliable a method of determining BSN175 stability. Therefore, IR should be the primary means of monitoring stability, although further research is necessary to build a comprehensive analytical package for confirming the identity and purity of BSN175.

Additionally, this research supports the use of Bootstrap Error-adjusted Single-sample Technique (BEST) statistical analysis, as its results are extremely clear and easy to interpret. The programs can also be implemented by anyone, despite their knowledge of computer programming, greatly increasing its number of applications.

## FUTURE RESEARCH

This is only one small piece of the work necessary to receive FDA approval for this drug. Although it seems definitive that IR spectroscopy is that best option for determining stability, this pilot research would need to be repeated in a cGLP environment. This experiment was simply preliminary work to understand how the drug decomposes and how to design a cGLP study.

Additional research would also need to be conducted to determine the best analytical test(s) for determining purity.

## ACKNOWLDEGMENTS

The project described was supported by the National Center for Research Resources and the National Center for Advancing Translational Sciences, National Institutes of Health, through Grant UL1TR001998, and through NSF ACI-1053575 allocation number BIO170011. The content is solely the responsibility of the authors and does not necessarily represent the official views of the NIH. A.A. thanks Dr. John Selegue for allowing access to his lab and supplying solvents and NMR tubes, and Dr. David Atwood for advice on troubleshooting the IR spectrometer.

## Appendix I Main Program

~~~
% Copyright 2003 Robert A. Lodder
function [sds,sdskew] = qb(tnspec,btrain,newspec,cnter,radfrac,sensitiv)
% function definition of qb with parameters
% tnspec= which is training spectra
% btrain =bootstrap replicates calculated using routine replica
% newspec= sample spectrum
% cnter= center of calibration set calculated using routine replica
% radfrac= fraction of points in hypercylinder
% sensitiv= sensitivity
b = size(btrain,1); %finds the number of rows in btrain
qdist = zeros(1,b); %creates a null row matrix
s02 = sqrt(sum((newspec-cnter).∧2)); %computing the squareroot of sum of the suares
s0r = sqrt(sum((btrain-repmat(cnter,b,1)).∧2,2)); %repmat creates a bX1 tilings of cnter
s2r = sqrt(sum((btrain-repmat(newspec,b,1)).∧2,2));
sub = (s02+s0r+s2r)/2;
area = sqrt(sub.*(sub-s02).*(sub-s0r).*(sub-s2r));
radial = (2*area)/s02;
project = sqrt(s0r.∧2-radial.∧2);
locs = find((s02.∧2+s0r.∧2) < s2r.∧2); %finds the indices where s02*s02 + s0r*s0r < s2r*s2r(getting the locus)
project(locs) = project(locs)*-1;
qdist = project;
qrr = sort(radial); %sorts the elements in ascending order
radii = qrr(radfrac*b);
qdist(find(radial > radii)) = 0; % setting the elments of qdist to zero where radial > radii
qdist = sort(qdist(find(qdist))); % sorts all non zero elememts in qdist
lindex = floor(0.16*length(qdist)); % lower limit found by rouding to nearest integer
uindex = floor(0.84*length(qdist)); % upper limit found by rounding to nearest integer
if(length(qdist) < 50)
 ’** Need more replicates in hypercylinder **’
end
sd = std(qdist)*sqrt(size(tnspec,1)); %std returns the standard deviation
sds = sqrt(sum((cnter-newspec).∧2))/sd; %calculation of the standard deviation distances
% BIAS ADJUSTMENT
alpha = normcdf(-1,0,1); % computes the cumulative distribution function with mean 0 and standard deviation 1
za = norminv(alpha,0,1); % inverse of the cumulative ditribution function with mean o and standard deviation 1
tcenter = median(tnspec); % finding the median
cs02 = s02;
cs0r = sqrt(sum((tcenter-cnter).∧2));
cs2r = sqrt(sum((tcenter-newspec).∧2));
csub = (cs02+cs0r+cs2r)/2;
carea = sqrt(csub*(csub-cs02)*(csub-cs0r)*(csub-cs2r));
cradial = (2*carea)/cs02;
cproject = sqrt(cs0r∧2-cradial∧2);
if((s02∧2+cs0r∧2) > cs2r∧2)
 cproject = −cproject;
end
n = length(qdist); % finds the length of the vector
if(floor(n/2) == n/2)
 md = (qdist(n/2)+qdist(n/2+1))/2;
else
 md = qdist(floor(n/2+0.5));
end
cproject = cproject*sensitiv + md;
fdist = qdist-cproject;
index = 1:length(fdist);
if(cproject > max(qdist))
 zelement = length(qdist)-1;
elseif(cproject < min(qdist))
 zelement = 1;
else
 rootloc = find(abs(fdist)==min(abs(fdist)));
 zelement = rootloc(1);
end
z0 = norminv(zelement/length(qdist),0,1);
if(abs(2*z0) > abs(za))
 error(’ –– Decrease skew sensitivity. ––’);
end
sensitiv = abs(sensitiv);
lowind = floor(normcdf(2*z0+za,0,1)*length(qdist));
upind = floor(normcdf(2*z0-za,0,1)*length(qdist));
if(lowind < 2)
 ’** Warning ** Too few replicates’
end
if(upind > length(qdist)-2)
 ’** Warning ** Too few replicates’
end
if(lowind < 1)
  lowind = 1;
end
if(upind > length(qdist))
  upind = length(qdist);
end
lowlim = qdist(lowind);
uplim = qdist(upind);
euc = sqrt(sum((cnter-newspec).∧2));
fac = abs(norminv(alpha));
erd = sqrt(size(tnspec,1));
if(abs(2*z0)>abs(fac))
  ’** Warning ** SKEW CORRECTION exceeds replicates’
end
sdskew = euc/((uplim/fac)*erd);
~~~

### Replica Program

~~~
function [BTRAIN,CNTER]=replica(TNSPEC,B)
% TNSPEC=training spectra, B=number of replicates desired.
% REPLICA outputs BTRAIN replicates, and the center of the replicates in CNTER
% “Copyright 2003 Robert A. Lodder & Bin Dai”
 [N,D]=size(TNSPEC);
 BTRAIN=zeros(B,D);
 CNTER=zeros(D,1);
 BSAMP=zeros(N,D);
 PICKS=rand(B*N,1);
 index=find(PICKS==1);
 PICKS(index)=0.9999;
 PICKS=reshape(PICKS,B,N);
 PICKS=fix(N*PICKS+1);
 for I=1:B
  BSAMP=TNSPEC(PICKS(I,:),:);
  BTRAIN(I,:)=sum(BSAMP)/N;
 end
 BTRAIN;
 CNTER=sum(BTRAIN)/B;
~~~

### Data Processing/Commands: [IR COMPARISONS]

~~~
>> IR_decomposed=zeros(2,2);
>> wavenumber=zeros(2,2);
>> plot(wavenumber,IR_decomposed);
>> plot(wavenumber,IR_decomposed);
>> IR_stable=zeros(2,2);
>> plot(wavenumber,IR_stable);
>> hold on
>> plot(wavenumber,IR_decomposed);
>> [BTRAIN,CNTER]=replica(IR_stable,1000);
>> [sds,sdskew]=qb(IR_stable,BTRAIN,IR_stable(1,:),CNTER,1,0);
>> [sds,sdskew]=qb(IR_stable,BTRAIN,IR_decomposed(1,:),CNTER,1,0);

<u>>> dist=mahal(IR_decomposed(1,:),IR_stable)</u>
<u>Error using mahal (line 38)</u>
<u>The number of rows of X must exceed the number of columns.</u>

>> sds
sds = **35.0483**
>>[sds,sdskew]=qb(IR_stable,BTRAIN,IR_decomposed(2,:),CNTER,1,0);
>> sds
sds = **24.7354**
>>[sds,sdskew]=qb(IR_stable,BTRAIN,IR_decomposed(3,:),CNTER,1,0);

>> sds
sds = **23.4584**
>>[sds,sdskew]=qb(IR_stable,BTRAIN,IR_decomposed(4,:),CNTER,1,0);
>> sds
sds = **22.1090**
>>[sds,sdskew]=qb(IR_stable,BTRAIN,IR_decomposed(5,:),CNTER,1,0);
>> sds
sds = **21.7209**
>>[sds,sdskew]=qb(IR_stable,BTRAIN,IR_decomposed(6,:),CNTER,1,0);
>> sds
sds = **25.7087**
~~~

### [IR CONTROLS: STABLE-STABLE]

~~~
>> [sds,sdskew] = qb(IR_stable,BTRAIN,IR_stable(1,:),CNTER,1,0);
>> sds
sds = **1.4076**
>> [sds,sdskew] = qb(IR_stable,BTRAIN,IR_stable(2,:),CNTER,1,0);
>> sds
sds = **0.7779**
>> [sds,sdskew] = qb(IR_stable,BTRAIN,IR_stable(3,:),CNTER,1,0);
>> sds
sds = **1.3360**
>> [sds,sdskew] = qb(IR_stable,BTRAIN,IR_stable(4,:),CNTER,1,0)
sds = **1.4371**
>> [sds,sdskew] = qb(IR_stable,BTRAIN,IR_stable(5,:),CNTER,1,0);
>> sds
sds = **1.4177**
>> [sds,sdskew] = qb(IR_stable,BTRAIN,IR_stable(6,:),CNTER,1,0);
>> sds
sds = **1.7298**
~~~

### [NMR COMPARISONS]

~~~
>> NMR_stable=zeros(2,2);
>> NMR_stable=transpose(NMR_stable);
>> PPM=zeros(2,2);
>> NMR_decomposed=zeros(2,2);
>>NMR_decomposed=transpose(NMR_decomposed);
>> plot(PPM,NMR_decomposed);
hold on
plot(PPM,NMR_stable);
>> hold off
>> plot(PPM,NMR_decomposed);
>> hold on
>> plot(PPM,NMR_stable);
>>[BTRAIN,CNTER]=replica(NMR_stable,1000);
>>[sds,sdskew]=qb(NMR_stable,BTRAIN,NMR_decomposed(1,:),CNTER,1,0);
>> sds
sds = **3.4954**
>>[sds,sdskew]=qb(NMR_stable,BTRAIN,NMR_decomposed(2,:),CNTER,1,0);
>> sds
sds = **3.6264**
>>[sds,sdskew]=qb(NMR_stable,BTRAIN,NMR_decomposed(3,:),CNTER,1,0);
>> sds
sds = **1.2395**
>>[sds,sdskew]=qb(NMR_stable,BTRAIN,NMR_decomposed(4,:),CNTER,1,0);
>> sds
sds = **1.1870**
>>[sds,sdskew]=qb(NMR_stable,BTRAIN,NMR_decomposed(5,:),CNTER,1,0);
>> sds
sds = **4.2819**
>>[sds,sdskew]=qb(NMR_stable,BTRAIN,NMR_decomposed(6,:),CNTER,1,0);
>> sds
sds = **4.3127**
~~~

### [NMR CONTROLS: STABLE-STABLE]

~~~
>> [sds,sdskew] = qb(NMR_stable,BTRAIN,NMR_stable(1,:),CNTER,1,0);
>> sds
sds = **1.2502**
>> [sds,sdskew] = qb(NMR_stable,BTRAIN,NMR_stable(2,:),CNTER,1,0);
>> sds
sds = **1.5016**
>> [sds,sdskew] = qb(NMR_stable,BTRAIN,NMR_stable(3,:),CNTER,1,0);
>> sds

sds = **0.7960**
>> [sds,sdskew] = qb(NMR_stable,BTRAIN,NMR_stable(4,:),CNTER,1,0);
>> sds
sds = **0.7391**
>> [sds,sdskew] = qb(NMR_stable,BTRAIN,NMR_stable(5,:),CNTER,1,0);
>> sds
sds = **1.0678**
>> [sds,sdskew] = qb(NMR_stable,BTRAIN,NMR_stable(6,:),CNTER,1,0);
>> sds
sds = **1.0260**
~~~

